# Olfactory decoding is positively associated with *ad libitum* food intake in sated humans

**DOI:** 10.1101/2022.06.01.494392

**Authors:** Emily E. Perszyk, Xue S. Davis, Dana M. Small

**Affiliations:** Modern Diet and Physiology Research Center, New Haven, CT, USA, 06510; Yale University School of Medicine, Department of Psychiatry, New Haven, CT, USA, 06510; Yale University, Department of Psychology, New Haven, CT, USA, 06510

**Keywords:** olfaction, intake, satiety, obesity, fMRI, multi-voxel pattern analysis

## Abstract

The role of olfaction in eating behavior and body weight regulation is controversial. Here we used functional magnetic resonance imaging to test whether central olfactory coding is associated with internal state, food intake, and change in body weight over one year in healthy human adults. Since odor quality and category are coded across distributed neural patterns that are not discernible with traditional univariate analyses, we used multi-voxel pattern analyses to decode patterns of brain activation to food versus nonfood odors. We found that decoding accuracies in the piriform cortex and amygdala were greater in the sated compared to hungry state. Sated decoding accuracies in these and other regions were also associated with post-scan *ad libitum* food intake, but not with weight change. These findings demonstrate that the fidelity of olfactory decoding is influenced by internal state and is associated with immediate food intake, but not longer-term body weight regulation.

## 1. Introduction

Olfaction is thought to play a critical role in energy consumption and body weight regulation. For instance, odors aid in localizing potential food sources, determining edibility, and preparing the body for subsequent ingestion (Stevenson, 2010). Odors can originate in the environment (e.g., smelling cookies in a bakery) and be sensed orthonasally through the nose, or they may emanate from in the oral cavity (e.g., taking a bite of cookie) and be sensed retronasally as flavor (Small and Prescott, 2005). Through both olfactory routes, food aromas stimulate salivation, gastric acid secretion, and insulin release (Lee and Linden, 1992; Feldman and Richardson, 1986; Yeomans, 2006; Proserpio et al., 2017). The smell of a food in the absence of visual or gustatory cues also transiently inhibits orexigenic agouti-related peptide (AgRP) neurons while activating anorexigenic proopiomelanocortin (POMC) neurons of the hypothalamus (Betley et al., 2015; Chen et al., 2015).

Despite these data showing that olfaction is integral to ingestive behavior, the exact nature of the relationship between olfactory function and obesity risk is controversial. On one hand, multiple lines of evidence point to a negative association between olfactory ability and both eating and body weight. People with olfactory deficits often report that they overeat to combat reduced food pleasure (Temmel et al., 2002; Aschenbrenner et al., 2008), have high Body Mass Index [BMI] (Richardson et al., 2004; Patel et al., 2015), and are susceptible to weight gain (Mattes et al., 1990; Mattes and Cowart, 1994). Accordingly, mice with genetically enhanced odor detection and discrimination are resistant to high fat diet-induced obesity (Xu et al., 2003; Fadool et al., 2004). On the other hand, separate work suggests a positive relationship, whereby olfactory loss is associated with decreases in food intake and body weight. Mice made anosmic by conditional ablation of olfactory sensory neurons are less vulnerable to weight gain and show blunted high-fat food consumption (Riera et al., 2017) compared to control animals. In humans, poor olfactory perception corresponds with lower BMI (Hubert et al., 1980; Stafford and Whittle, 2015) and undereating in advanced age that can be restored by amplifying food flavoring (Schiffman, 1998).

It is also well-recognized that the nutritional status of an organism can influence odor perception. Starvation increases olfactory discrimination in *C. elegans* (Colbert and Bargmann, 1997), sensitizes olfactory sensory neurons in *Drosophila* (Root et al., 2011), and enhances mitral cell food odor reactivity (Pager et al., 1972), olfactory bulb Fos expression (Prud’homme et al., 2009), and the ability to detect minute concentrations of aversive odorants in rodents (Aimé et al., 2007). Cells of the olfactory bulb express receptors for a number of satiety-regulating hormones (Hill et al., 1986; Gall et al., 1986; Elmquist et al., 1998; Thiebaud et al., 2016; Palouzier-Paulignan et al., 2012; McIntyre et al., 2017). In turn, such hormones as ghrelin and leptin are capable of influencing olfactory acuity in a way that mimics the effects of food deprivation and consumption (Aimé et al., 2007; Julliard et al., 2007; Tong et al., 2011; Cameron et al., 2012). In humans, the relationship between internal state and odor acuity may be impacted by adiposity and ghrelin sensitivity (Sun et al., 2016). However, there is strong evidence that odor-evoked brain responses decrease in parallel with satiety (Boesveldt, 2017; Gervais and Pager 1979, O’Doherty et al., 2000; Prud’homee et al., 2009 and Soria Gomez et al., 2014) and that those specific to food odors correlate with BMI change (Sun et al., 2015; Han et al., 2020).

There is also evidence that odor quality (e.g., rose versus lemon) and category (e.g., food versus nonfood) are encoded in distributed neural patterns within primary olfactory cortex (Howard et al., 2009; Stettler and Axel, 2009; Qu et al., 2016; Bao et al., 2016) and that this coding is influenced by internal state. Specifically, functional neuroimaging studies of olfaction have employed multi-voxel pattern analyses (MVPA) to interrogate odorant-specific activity patterns and report evidence that the accuracy of pattern decoding is influenced by sleep deprivation (Bhutani et al., 2019) and food consumption (Shanahan et al., 2021). Whether and how this translates to eating behavior and weight gain susceptibility is unclear and constitutes an important gap because food cue reactivity is a prominent obesity risk factor (Boswell and Kober, 2016). Exposure to food odors leads to increased eating and craving of cued foods (Gaillet et al., 2013), especially in restrained eaters (Fedoroff et al. 1997; Fedoroff et al. 2003).

Here we sought to test if the fidelity of olfactory decoding and/or averaged odor-evoked responses in the piriform cortex and amygdala are (1) modulated by nutritional status and (2) related to food intake or weight change over one year. To this end, we re-analyzed a prior dataset in which participants were presented with orthonasal food and floral (nonfood) odors during functional magnetic resonance imaging (fMRI) in hungry and sated states. Our previous univariate analyses of this data showed that averaged responses to food odor presentations in the amygdala, but not piriform cortex, were negatively associated with weight change only in the hungry state (Sun et al., 2015). However, MVPA was not used, raising the possibility that these analyses were not sensitive to odor coding. Further, responses to the two odorant categories were not contrasted directly. In the current study, we therefore used MVPA to quantify fMRI odorant category decoding and univariate analyses to capture average activation for food versus nonfood odors. We then compared these measures to post-scan *ad libitum* intake and weight change over one year. We reasoned that if heightened olfactory function confers risk for obesity, then better decoding accuracy of food and nonfood odors in olfactory cortex should positively correlate with food intake and weight change. Alternatively, if worse olfactory function leads to greater obesity risk, then negative relationships should emerge. We also tested whether these effects were specific to olfactory decoding or generalized to the magnitude of univariate responses.

## 2. Methods

### 2.1 Participants

The current work is a reanalysis of a previously acquired and published dataset (Sun et al., 2016, 2015). Thirty-three healthy right-handed participants [17 male, 16 female; age: 27.0 ± 6.3 (18−40) years; body mass index (BMI): 24.01 ± 3.87 (19.50−33.60) kg/m^2^; M ± SD (range)] took part in the study. Seventeen identified as White, eight as Asian, six as Black or African American, and two did not report their race. Additionally, three reported being Hispanic/Latinx, 26 non-Hispanic/Latinx, and four did not report their ethnicity. Participants were recruited from the greater New Haven, Connecticut area through the Yale University Interdisciplinary Research Consortium on Stress, Self-Control and Addiction (IRCSSA) P30 Subject’s core and via flyer advertisements. Prior to study enrollment, participants were screened to be free of chemosensory impairments, neurologic or psychiatric disorders, head injury with loss of consciousness, lactose intolerance, and food allergies. Participants could not be pregnant or nursing, currently dieting, or using tobacco or drugs other than alcohol. Females completed their scans on days five to nine or 15 to 28 of their self-reported 28-day menstruation cycle to avoid scanning during menstruation or ovulation. All participants provided written informed consent and the study procedures were approved by the Yale Human Investigations Committee.

### 2.2 Stimuli and delivery

#### 2.2.1 Odors

The food odors included chocolate (chocolate cookie, 6002335, Bell Flavors and Fragrances, Northbrook, IL, USA) and strawberry (strawberry and cream, 6106524, Bell Flavors and Fragrances). The nonfood odors were honeysuckle (Chey N-3, 039831, Firmenich SA, Geneva, Switzerland) and lilac (lilac 71, 31731066, International Flavors and Fragrances, New York, NY, USA). Odors were delivered with a custom MRI-compatible olfactometer programmed in Labview (National Instruments, Austin, TX, USA) that has been previously described (Small et al., 2008). In short, the flow of humidified and temperature-controlled air was adjusted by mass flow controllers (MKS Instruments, Andover, MA, USA). This allowed clean air to pick up vaporized odor molecules by passing over stainless steel odor-containing wells. Odorized air from separate channels converged in a steel mixing manifold and exited through one of two Teflon tubes, the first of which was dedicated to odors and the second to clean air. These tubes were each coupled to a vacuum line through a Teflon manifold (Teqcom, Santa Ana, CA, USA) that rested on the participant’s chest. The vacuums allowed for a closed-loop system that removed air to prevent head space contamination. Participants wore a nasal mask (Philips Respironics, Murrysville, PA, USA) through which odor stimuli or clean air entered in a continuous stream from the manifold and exited by vacuum. Finally, the nasal mask was attached to a pneumotachograph that measured airflow in the nose (Johnson and Sobel, 2007) and subsequently to a spirometer (ADInstruments, Sydney, Australia) that provided a signal which was amplified (ADInstruments, PowerLab 4SP) and digitally recorded at 100 Hz with Chart 5.5.6 (ADInstruments). Each odor was diluted with clean air to achieve equivalent ratings of moderate intensity across all participants during a separate training session. Odor dilutions were held constant within participants across all sessions.

#### 2.2.2 Standardized meals

For breakfast, participants received Nature’s Valley brand Crunchy variety granola bars. Lunches consisted of apple slices (∼25 kcal per serving) and the choice of tuna, ham, turkey, or avocado sandwiches prepared with white bread, Kraft American cheese, tomato, and mayonnaise (∼400 kcal per sandwich).

#### 2.2.3 Ad libitum foods

Two flavors of milkshakes (chocolate and strawberry) in opaque cups and a large tub of Annie’s brand Shells and White Cheddar (cheese pasta) totaling ∼1750 kcal were used for the post-scan measure of *ad libitum* intake. The chocolate milkshake was a combination of 12 fl oz whole milk, Garelick Farms brand Chug Chocolate Milkshake, and Garelick Farms brand Chug Cookies and Cream Milkshake. The strawberry milkshake consisted of 32 fl oz whole milk and 6 fl oz Hershey’s brand strawberry syrup.

### 2.3 Experimental procedures

Participants completed six total sessions. The first five sessions occurred on separate days within three months and included one training session, one behavioral session, and three fMRI scanning sessions (hungry, sated, and *ad libitum* conditions). The final session was a one-year follow-up. Only data from the hungry and sated scan conditions [elapsed days between sessions: 22.8 ± 14.6 (7–70); M ± SD (range)] were re-analyzed here. This was because of extreme variability in pre-scan energy intake during the *ad libitum* condition that could have confounded the results.

#### 2.3.1 Training and behavioral sessions

Participants were instructed to arrive to the training and behavioral sessions neither hungry nor full but fasted for at least one hour from any foods or drinks besides water. The participant’s height was measured with a digital stadiometer and weight with an electronic scale. BMI in kg/m^2^ was calculated as weight divided by the square of height. Participants completed the Dutch Eating Behavior Questionnaire (DEBQ), which includes scales for restrained, emotional, and external eating behaviors (Strien et al., 1986). They were also trained to make computerized internal state and stimulus ratings. Internal state ratings including hunger, fullness, thirst, anxiety, and need to urinate were made on adapted 100mm cross-modal general labeled magnitude scales (gLMS) from “barely detectable” to “strongest imaginable sensation” (Bartoshuk et al., 2004; Green et al., 1996, 1993). Stimulus ratings included intensity, liking, familiarity, wanting to eat, and edibility. Intensity and liking were rated on 100mm category-ratio scales: the gLMS (as above), and the Labeled Hedonic Scale (Lim et al., 2009) from “most disliked” to “most liked sensation imaginable,” respectively. Familiarity, wanting to eat, and edibility were rated on 200mm horizontal visual analog scales labeled at the left (–100) anchor, center (0) with “neutral,” and right (+100) anchor. The anchor labels for familiarity were “not familiar at all” (–100) and “very familiar” (+100), for wanting to eat were “I would never want to consume this” (–100) and “I would want to consume this more than anything” (+100), and for edibility were “not edible at all” (–100) and “very edible” (+100).

Finally, participants were brought to a mock fMRI scanner that was set-up with the gustometer and olfactometer systems. Participants were first exposed to each odor separately, and an experimenter manually adjusted the odor concentrations to their verbal ratings of moderate intensity. Participants also practiced making internal state and stimulus ratings after the delivery of each odor and flavor using a computer mouse and computer monitor back-projected to a headcoil-mounted mirror. Participants then entered the mock scanner bore and took part in simulations of one odor and one flavor run, after which they left the bore and completed a second round of internal state and stimulus ratings. Breakfast bars were given to each participant to take home and save for the morning of their scan sessions.

#### 2.3.2 fMRI scanning sessions

Prior to arriving for the scan at 11:30 AM, participants were instructed to eat the breakfast bars (one package for females, one and a half packages for males) at any time in the morning and then to refrain from further eating or drinking besides water. Participants filled out paperwork at the start of the session, including a food diary for that morning and the previous day. In cases where participants reported eating a breakfast other than the bars (or no meal) before their first scan, they were told to eat the same breakfast (or lack thereof) for the remaining scans to match morning intake across sessions. At ∼12:50 PM, participants completed the lunch manipulation. They consumed either a fixed-portion meal that was adjusted by sex to constitute approximately 25% of daily energy need (Sated condition, consisting of one sandwich and one serving of apple slices for females, one and a half sandwiches and one serving of apple slices for males) or nothing (Hungry condition). Except for a participant who removed the cheese from the sandwich and another who didn’t eat the apple slice skins, participants ate the entire meal.

The experimental design for each scan is summarized in **Figure 1a**. Prior to the meal (or no meal), a Teflon catheter was inserted into an antecubital vein for blood sampling. Participants rated their internal state twice concomitant with two IV blood draws before and after eating the standardized lunch (or no meal) and were then inserted into the scanner bore. Prior to scanning, stimulus ratings for each odor were assessed. Participants completed four odor and two flavor runs per scan with their eyes closed. Each odor run lasted ∼6min and contained six food odor, six nonfood odor, and six clean air trials. Trials began with an auditory cue of “three, two, one, sniff.” Odor or clean air delivery was time-locked to sniff onset and lasted for 3s followed by a 9–19s intertrial interval. The flavor runs have been detailed in prior publications (Sun et al., 2015, 2014). Internal state ratings and IV blood draws were collected again at 30, 60, and 90min from the meal (or no meal) onset, and final stimulus ratings were made at 90min before participants were removed from the scanner.

**Figure 1.**
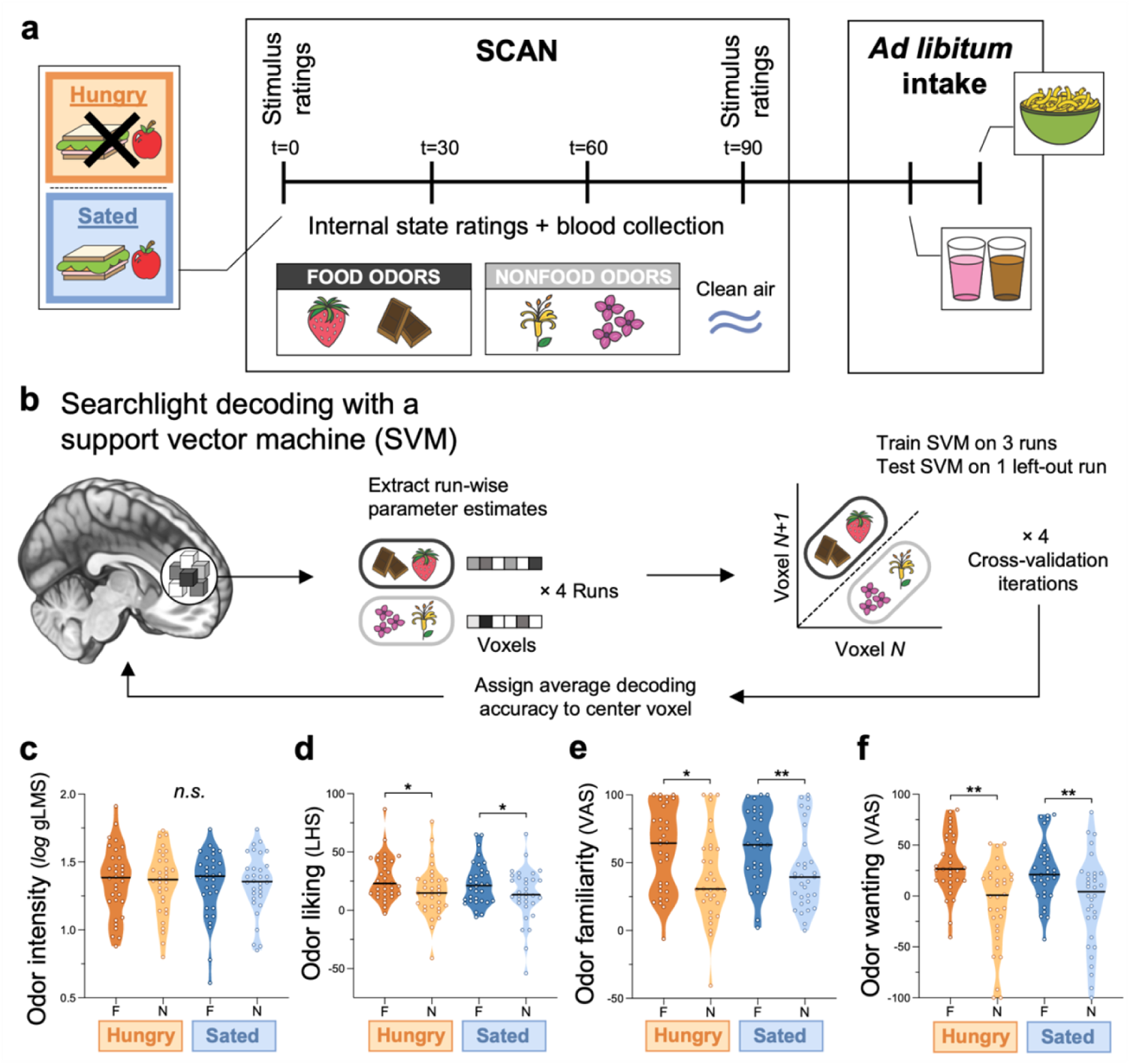
Experimental design and behavioral outcomes. **(a)** Session overview for the fMRI scans completed when participants were hungry (no meal) or sated after a fixed-portion meal. Time refers to minutes post-meal (or no meal). For each scan, food (strawberry and chocolate) and nonfood (honeysuckle and lilac) odors and clean air were presented in pseudorandomized order during 4 runs of fMRI scanning. Participants were instructed to consume strawberry and chocolate milkshake and cheese pasta *ad-libitum* after the scan. **(b)** Schematic of the whole-brain searchlight multi-voxel pattern analyses in which support vector machines were used to decode food from nonfood odors using their run-wise voxel parameter estimates. Truncated violin plots show differences in perceptual ratings of odor **(c)** intensity, **(d)** liking, **(e)** familiarity, and **(f)** wanting to eat when participants were hungry versus sated. Each data point depicts a single participant and shading represents the density of the points around the median line in black. T, time; F, food odors; N, nonfood odors; *n.s.*, not significant; *\* p* < 0.001; *\*\* p* < 0.0001.

After the scan, participants were brought to a separate room where they were offered the opportunity to eat as much of the *ad libitum* foods (chocolate and strawberry milkshake and pasta) as they liked. Participants were unaware that the weights of the milkshakes and pasta were recorded in grams before and after consumption to quantify intake.

#### 2.3.3 fMRI scanner

Imaging data were acquired on a Siemens 3T TIM Trio Scanner at the Yale University Magnetic Resonance Research Center. High resolution T1-weighted structural images were acquired for each participant with repetition time (TR) = 2230ms, echo time (TE) = 1.73ms, flip angle = 9°, matrix = 256×256, slice thickness = 1mm, field of view (FOV) = 250×250, 176 slices. A susceptibility-weighted single-shot echoplanar sequence was used to capture regional distribution of the blood-oxygen-level-dependent (BOLD) signal. The signal was allowed to equilibrate over six scans (12s) at the start of each functional run, which were discarded prior to the analyses. The functional images were acquired with TR = 2000ms, TE = 20ms, flip angle = 80°, FOV = 220, matrix = 64×64, slice thickness = 3mm. Forty contiguous slices were collected in an interleaved method to reduce the crosstalk of the slice section pulse.

#### 2.3.4 One-year follow-up session

Participants returned to the lab as close as possible to one year from the date of their behavioral session [elapsed weeks: 53.0 ± 3.0 (49.1−61.3) weeks; M ± SD (range)]. Follow-up BMI was measured as before and change in BMI (ΔBMI) was calculated. Two participants moved away from New Haven and were unable to return. They were instructed to weigh themselves on their own digital scale with minimal clothing and self-report their new weight. Four other participants neither returned nor self-reported and were excluded from the analyses on weight change.

### 2.4 Data analyses

#### 2.4.1 Behavioral analyses

The behavioral data were analyzed in MATLAB R2020a (MathWorks, Natick, MA, USA) and plotted using GraphPad Prism version 9.2.0 (GraphPad Software, San Diego, CA, USA). Paired Student’s *t*-tests were used to examine differences in satiation index and *ad libitum* intake in the fed versus fasted state. To determine if perceptual ratings varied as a function of internal state or odor category, a linear mixed model was conducted for each with internal state (hungry/sated), odor category (food/nonfood), and the interaction of the two as fixed effects, and participant as a random effect. Linear regressions were performed to identify measures associated with food intake and weight change. The goal of these steps was to determine variables that should be included as covariates in the fMRI analyses (e.g., odor ratings, BMI, and satiation index), so correction for multiple comparisons was not performed. Prior to formal analyses, each participant internal state and odor intensity rating made on the gLMS was first log_10_ transformed. The four internal state ratings from timepoints 0, 30, 60, and 90 were then averaged, and a satiation index was calculated by subtracting hunger from fullness within participant and scan condition. Pre-and post-scan ratings of odor intensity, liking, familiarity, wanting to eat, and edibility were also averaged separately for the food and nonfood odor categories. The effect of time on these ratings was reported previously (Sun et al., 2016). Odor ratings for wanting to eat and edibility were highly correlated when hungry (*r^2^* = 0.796, *p* < 0.00001) and when sated (*r^2^* = 0.754, *p* < 0.00001). The current analyses focus on wanting to eat to eliminate redundancy in this construct; edibility ratings of the food and nonfood odors have also been published in the past (Sun et al., 2016). Internal state and odor ratings were missing for one participant due to technical malfunction, leaving n = 32 for analyses with these variables.

Post-scan *ad libitum* milkshake and pasta intake were converted from grams to kilocalories using the nutritional information provided on the package labels and then summed. Spirometer data were analyzed in Chart 5.5.6 (ADInstruments) as described in prior work (Sun et al., 2016). In brief, the start and end of each sniff were defined as the time of odor onset and the trough when airflow reached a minimum prior to the next inhalation, respectively. Sniff volume and vigor were calculated for all odors per participant and scan condition over 3 steps. First, for each individual sniff, sniff volume was measured as the area under the curve (minus baseline), and sniff vigor was quantified as the maximum slope. Second, sniff volume and vigor were averaged in response to all odor presentations during each scan. Third, sniff volume and vigor were separately divided by the average volume and vigor of 20 randomly selected non-sniff breaths from the same scan condition to account for variation in breathing at rest.

Spirometer data from 12 participants were unusable due to technical issues during scanning; therefore, n = 21 for analyses with sniff vigor and volume included as covariates.

#### 2.4.2 Ghrelin analyses

Collected blood samples were immediately centrifuged post-scan, kept on ice until the completion of the session, and then stored at –80°C. Total circulating ghrelin levels were assessed per sample by the double antibody technique with a commercially-available radioimmunoassay (catalog #GHRT-89HK; MilliporeSigma, Burlington, MA, USA) using ^125^I-labeled ghrelin with ghrelin antiserum. For the hungry scan, circulating ghrelin level was quantified as average total ghrelin across all time points since no meal was consumed. In contrast, maximal plasma ghrelin change from time 0 to 30, 60, or 90 minutes was used for the sated scan to capture peak meal-related ghrelin excursion, in line with our previous plasma analyses (Sun et al., 2015). Four participants in the fasted scan and five in the fed scan were unable to tolerate blood sampling or their blood quantities collected were insufficient for biochemical measurement. Therefore, n = 29 for analyses on total average ghrelin when hungry and n = 28 for those on peak ghrelin excursion when sated, respectively.

#### 2.4.3 fMRI preprocessing

Neuroimaging data were preprocessed and analyzed on Linux workstations using MATLAB R2011a (MathWorks) and SPM12 (Statistical Parametric Mapping, Wellcome Centre for Human Neuroimaging, London, UK). Functional echo-planar images (EPIs) were time-acquisition corrected to the slice obtained at half of the TR, realigned to the mean, and co-registered to the average T1-weighted image. The anatomical T1 image was spatially normalized (1mm^3^ voxel size) to the Montreal Neurological Institute (MNI) template using the 6-tissue probability map from SPM12, resulting in deformation fields to transform the functional data (3mm^3^ voxel size). Prior to the univariate and connectivity analyses, these deformation fields were applied to the realigned and co-registered EPIs, which were then detrended (Macey et al., 2004) and smoothed using a 6mm full-width at half-maximum (FWHM) isotropic Gaussian kernel. For MVPA, the realigned and co-registered EPIs were left in the participant’s native space and detrended, but not smoothed (see below). The Artifact Detection Tools (ART) toolbox (Gabrieli Laboratory, McGovern Institute for Brain Research, Cambrige, MA, USA) was used to detect global mean and motion outliers in all EPI data. Motion parameters were included as regressors in the single-subject design matrix. Image volumes in which the z-normalized global brain activation exceed three SDs from the mean of the run or had more than 1mm of composite (linear plus rotational) voxel displacement were also flagged as outliers and de-weighted during SPM estimation. In addition, composite head motion per participant and scan was calculated as the sum of the average run-wise (linear plus rotational) voxel displacement across the four odor runs. It was used as a regressor at the group-level to further rule out any effects of motion.

#### 2.4.4 fMRI multi-voxel pattern analyses

To quantify olfactory decoding, searchlight-based MVPA was used. Food and nonfood odor categories were decoded from their underlying neural activity patterns separately for the hungry and sated scan conditions (**Fig. 1b**). First, whole-brain BOLD responses were estimated with separate general linear models (GLMs) for each participant and run using the non-normalized and non-smoothed EPI data. The (1) food odor, (2) nonfood odor, and (3) clean air presentations were modeled with a canonical hemodynamic response function (HRF) as events of interest with durations of 3s. The auditory sniff cue was modeled as an event of no interest, and the motion parameters were included as nuisance regressors. A 128s high-pass filter was applied to the time-series data to remove low-frequency noise and slow signal drifts. From this GLM, voxel-wise patterns were created for sniffing the food and nonfood odors by extracting their parameter estimates and subtracting the mean activity across the conditions in each run. In a leave-one-run-out, cross-validated approach, MVPA was performed on these voxel patterns localized to a moving spherical searchlight (3-voxel radius) using The Decoding Toolbox (Hebart et al., 2015). A support vector machine (SVM) implemented with the Library for Support Vector Machines (LIBSVM) package (Chang and Lin, 2011) was trained to decode food from nonfood odors using patterns of BOLD activation in three of the four runs. The SVM was then tested for its accuracy to predict these odor categories (food/nonfood) from the patterns in the left-out run. The average of the run-wise decoding accuracies across the four cross-validation iterations was mapped to the center voxel of each searchlight. The resulting whole-brain searchlight maps of decoding accuracy were normalized to MNI space using the participant’s deformation fields (see *fMRI preprocessing*) and smoothed (6mm FWHM).

To test where in the brain olfactory decoding is significantly greater than chance for each scan condition, the normalized and smoothed searchlight maps were subjected to random effects one-sample *t*-tests in SPM12. Sated > hungry decoding accuracies were also compared with a paired samples *t*-test. To assess the relationships between odor category decoding and food intake or weight change, total *ad libitum* energy and ΔBMI were regressed against whole-brain decoding accuracy in SPM12. Since BMI and satiation index were significantly correlated with food intake in the behavioral data, these variables were included as covariates in the brain regressions for intake, along with sex and age which are known to contribute to eating behavior (Martí-Henneberg et al., 1999). To determine if any relationships observed between decoding accuracies and food intake or ΔBMI could be attributed to the odor perceptual ratings, odor dilutions, composite head motion in the scanner, sniff vigor or volume, dietary restraint (quantified as the DEBQ restrained eating score), or circulating ghrelin levels, regressions between whole-brain decoding accuracies and these potential confounding variables were also performed.

#### 2.4.5 fMRI univariate analyses

Traditional univariate analyses were performed in SPM12 using the normalized and smoothed EPI data. The GLMs were the same as above (see *fMRI multi-voxel pattern analyses*). A two-way analysis of variance (ANOVA) was conducted on the contrasts for food odors > clean air and nonfood odors > clean air from the separate scan conditions to test for main effects or interactions of odor category or internal state on BOLD responses. *Ad libitum* intake (with covariates for BMI, sex, age, and satiation index) and ΔBMI were also regressed against univariate responses in the contrast of food > nonfood odors.

#### 2.4.6 fMRI connectivity analyses

A psychophysiological interaction (PPI) analysis (Friston et al., 1997) was carried out with the gPPI toolbox (McLaren et al., 2012) in SPM12 to identify the role of functional connectivity differences in the relationship uncovered between post-scan food intake and olfactory decoding in the amygdala when sated. The first eigenvariate of the time-series EPI data was extracted from the significant bilateral amygdala clusters that survived small volume correction (SVC) in the regression of sated decoding accuracy and adjusted food intake. BOLD activity in this amygdala seed therefore served as the physiological factor for the PPI model. PPI terms were created by deconvolving the eigenvariate (Gitelman et al., 2003), multiplying it with the psychological variable (food > nonfood odors), and reconvolving it with the HRF. New parameter estimates were computed for each participant by including the PPI term as a regressor of interest and the eigenvariate and psychological variables as nuisance regressors. Finally, decoding accuracy (from the peak voxel of the seed) and food intake were separately regressed against the PPI parameter estimate images for all participants in SPM12. This revealed brain areas in which food > nonfood connectivity with the seed region described variance in post-scan energy consumption or the fidelity of olfactory decoding in that seed.

#### 2.4.7 fMRI statistical analyses

Statistical thresholds were set to *p*_uncorrected_ < 0.001 and a cluster size of at least five contiguous voxels for all fMRI analyses. Both whole-brain and region of interest (ROI) analyses were performed with the univariate and MVPA approaches. The ROIs included (1) the piriform cortex created from the combination of olfactory cortex masks parcellated functionally (Zhou et al., 2019) and anatomically in the Automated Anatomical Labeling (AAL) atlas 3 (Rolls et al., 2020), and (2) the amygdala using the AAL atlas 3. Effects in these predicted ROIs were considered significant at a peak-level of *p* < 0.025, family-wise error small-volume corrected (*p*_FWE_) for multiple comparisons in the ROI masks and subsequently Bonferroni corrected for the two ROI searches. Unpredicted results were considered significant at *p* < 0.05, cluster-level family-wise error (*p*_FWE-cluster_) corrected across the whole brain.

## 3. Results

### 3.1 Behavioral results

As previously reported, ratings of satiety were greater and post-scan *ad libitum* energy intake was reduced after participants consumed a fixed-portion meal compared to when they were in a fasted state (Sun et al., 2016, 2015). We have also previously described the impact of internal state on perceptual ratings collapsed across odor category (Sun et al., 2016) along with the associations between these ratings and obesity risk (Sun et al., 2015). Here we adapt these analyses to compare the food versus nonfood odors; the results are reported in **Table 1** and **Fig. 1c-f**. Finally, our prior publication details the behavioral measures associated with weight change (Sun et al., 2015). Here we test the relationships between these measures and food intake to identify covariates for the subsequent fMRI analyses; the results are provided in **Table 2**. In particular, *ad libitum* intake positively correlated with participants’ subjective satiation when both fasted and fed, and with BMI when fed but not fasted.

**Table 1.** Perceptual ratings and their relationships with internal state, odor category, ad libitum food intake, and weight change.

| Perceptual rating | Linear mixed models |  |  | HUNGRY |  | SATED |  |
| --- | --- | --- | --- | --- | --- | --- | --- |
|  | Main effect of internal state | Main effect of odor category | Interaction | Intake | ΔBMI | Intake | ΔBMI |
| Intensity | $F_{(1,124)} = 0.311$,<br>$p = 0.578$ | $F_{(1,124)} = 0.160$,<br>$p = 0.690$ | $F_{(1,124)} = 0.114$,<br>$p = 0.736$ | $r^2 = 0.047$,<br>$p = 0.235$ | $r^2 = 0.075$,<br>$p = 0.152$ | $r^2 = 0.070$,<br>$p = 0.145$ | $r^2 < 0.001$,<br>$p = 0.973$ |
| Liking | $F_{(1,124)} = 2.375$,<br>$p = 0.126$ | $F_{(1,124)} = 15.180$,<br>**$p < 0.001^*$** | $F_{(1,124)} = 0.299$,<br>$p = 0.586$ | $r^2 = 0.005$,<br>$p = 0.690$ | $r^2 = 0.015$,<br>$p = 0.532$ | $r^2 = 0.074$,<br>$p = 0.133$ | $r^2 < 0.001$,<br>$p = 0.990$ |
| Familiarity | $F_{(1,124)} = 0.139$,<br>$p = 0.710$ | $F_{(1,124)} = 18.710$,<br>**$p < 0.001^*$** | $F_{(1,124)} = 0.154$,<br>$p = 0.696$ | $r^2 = 0.105$,<br>$p = 0.070$ | $r^2 = 0.028$,<br>$p = 0.390$ | $r^2 = 0.009$,<br>$p = 0.601$ | $r^2 < 0.001$,<br>$p = 0.936$ |
| Wanting to eat | $F_{(1,124)} = 1.326$,<br>$p = 0.252$ | $F_{(1,124)} = 36.477$,<br>**$p < 0.001^*$** | $F_{(1,124)} = 1.426$,<br>$p = 0.235$ | $r^2 = 0.012$,<br>$p = 0.551$ | $r^2 = 0.016$,<br>$p = 0.513$ | $r^2 = 0.021$,<br>$p = 0.428$ | $r^2 = 0.002$,<br>$p = 0.822$ |

\* Bold font denotes significant effects.

**Table 2.** Behavioral variables and their associations with ad libitum food intake.

| Behavioral variable | Intake |  |
| --- | --- | --- |
|  | HUNGRY | SATED |
| Subjective satiation | $r^2 = 0.155$,<br>**$p = 0.023^*$** | $r^2 = 0.238$,<br>**$p = 0.004^*$** |
| Dietary restraint | $r^2 = 0.043$,<br>$p = 0.248$ | $r^2 = 0.039$,<br>$p = 0.268$ |
| Ghrelin <sup>^</sup> | $r^2 = 0.028$,<br>$p = 0.387$ | $r^2 = 0.027$,<br>$p = 0.402$ |
| BMI | $r^2 = 0.078$,<br>$p = 0.117$ | $r^2 = 0.166$,<br>**$p = 0.019^*$** |

\* Bold font denotes significant effects; <sup>^</sup> Reflects total average ghrelin in the hungry state and peak ghrelin excursion in the sated state.

### 3.2 Imaging results

#### 3.2.1 Meal consumption enhances olfactory decoding

We first sought to uncover the impact of internal state on odor category decoding assessed with searchlight-based MVPA. **Table 3** outlines significant regions that emerged from the whole-brain and ROI analyses. The neural patterns activated while sniffing food versus nonfood odors were not discriminable in the hungry state. In contrast, the SVM could reliably classify food from nonfood odors bilaterally in the piriform cortex and in the amygdala (extending into the left uncus and superior temporal gyrus) when participants were sated (peak accuracies: left, 67.4%; right, 64.9%; chance-level = 50%). Since odor liking, familiarity, and wanting to eat were significantly greater for the food versus nonfood odors **(Table 1)**, we also included these perceptual ratings as covariates. Once again, the SVM decoding accuracy was greater than chance in the bilateral piriform cortex and in the amygdala only when sated. Significant decoding was also observed in the left superior temporal gyrus, inferior orbitofrontal gyrus, and uncus. We further compared sated > hungry decoding accuracy and observed positive effects in the bilateral amygdala and in the right piriform cortex (**Fig. 2a**). A cluster was observed in the left piriform cortex that did not survive correction for the two ROI searches (*t*_(32)_ = 3.695, *p*_FWE-cluster_ = 0.039, size = 12 voxels, x = –24, y = 5, z = −16). The other responses remained significant after covarying out the odor perceptual ratings, indicating that the increased decoding accuracies in the sated state were unlikely due to an effect of satiety on the value of the food versus nonfood odors. Interestingly, inclusion of the odor ratings in the model also resulted in significant decoding accuracy post-meal in the midbrain spanning both the substantia nigra and the ventral tegmental area (**Fig. 2b**).

**Figure 2.**
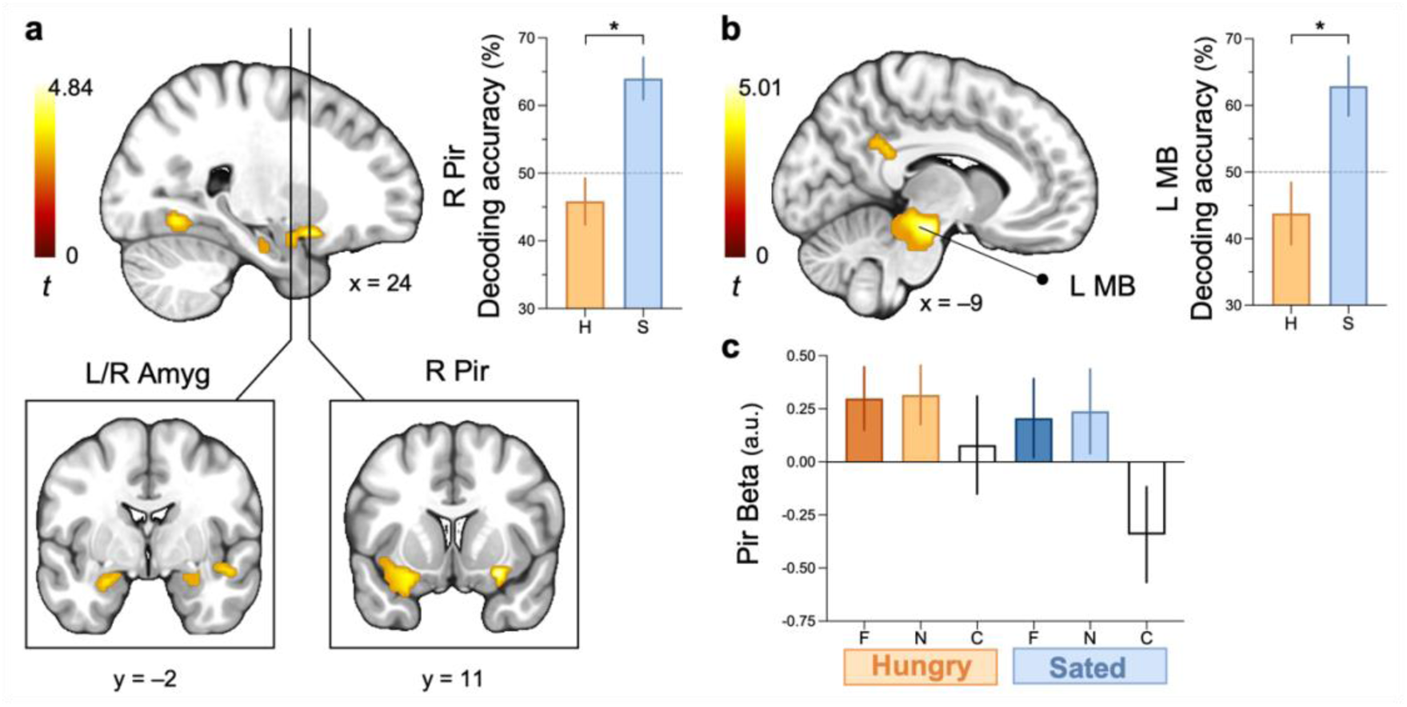
Effects of internal state on olfactory decoding and univariate responses to odors. SVM decoding accuracies for food versus nonfood odors were significantly greater when sated > hungry in **(a)** the right piriform cortex and the bilateral amygdala, and **(b)** the substantia nigra and ventral tegmental area of the left midbrain after adjusting for perceptual ratings of odor liking, familiarity, and wanting to eat. **(c)** A 2-way ANOVA on the univariate contrasts for food odors > clean air and nonfood odors > clean air by scan revealed no main effects or interaction of odor category or internal state on BOLD responses. Brain sections show the SPM *t*-map (*p*_uncorrected_ < 0.005, clusters of at least 5 voxels) overlaid onto an anatomical template in MNI coordinates. Decoding accuracies were extracted from the peak voxels and mean univariate beta estimates (from individual contrasts of food odors, nonfood odors, and clean air) were extracted from the piriform cortex ROI for illustrative purposes. Data are represented in bar plots as M ± SEM. L, left; R, right; Amyg, amygdala; MB, midbrain; Pir, piriform; H, hungry; S, sated; F, food odors; N, nonfood odors; C, clean air; a.u., arbitrary units; \**p*_FWE_ < 0.025.

**Table 3.**
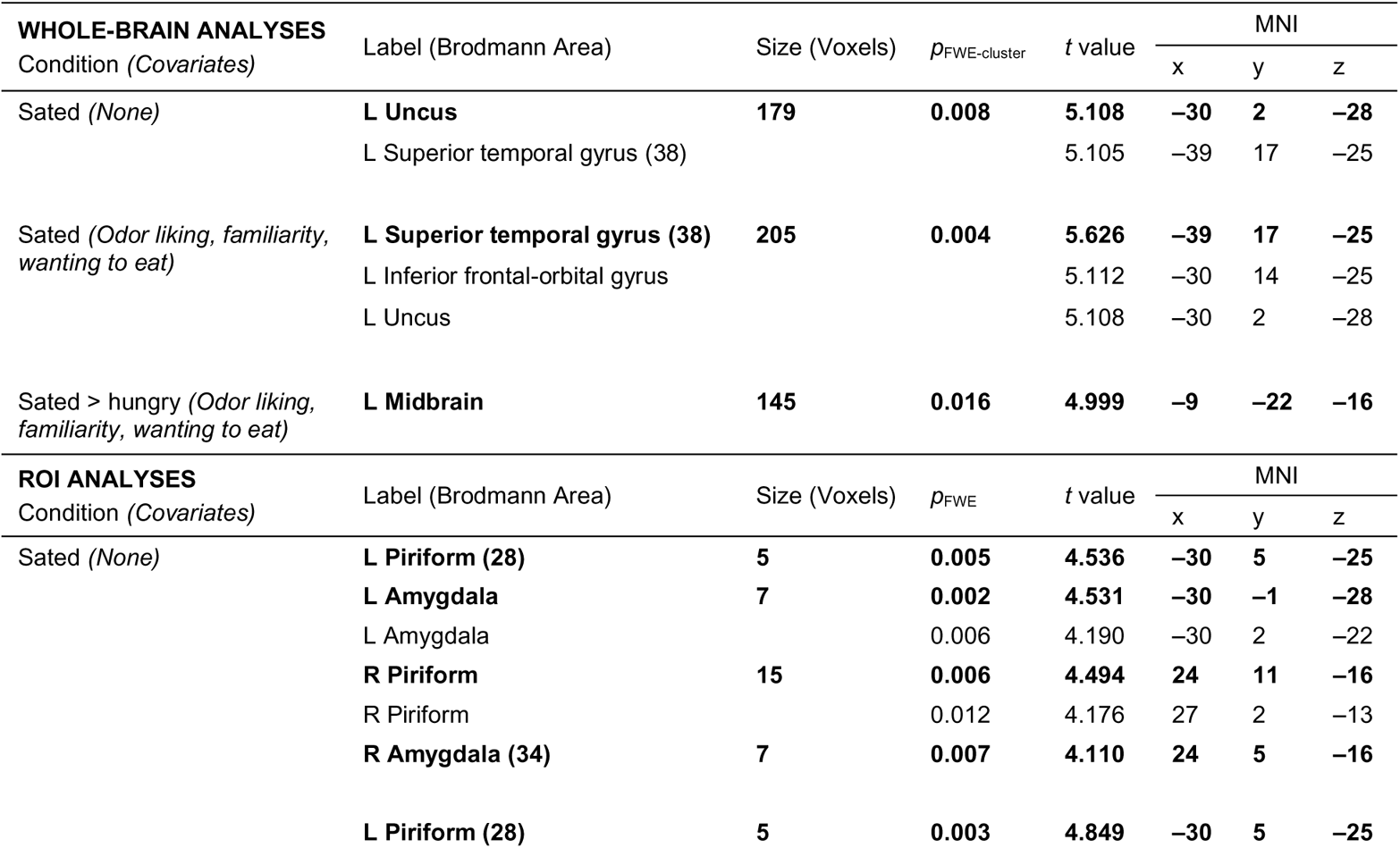
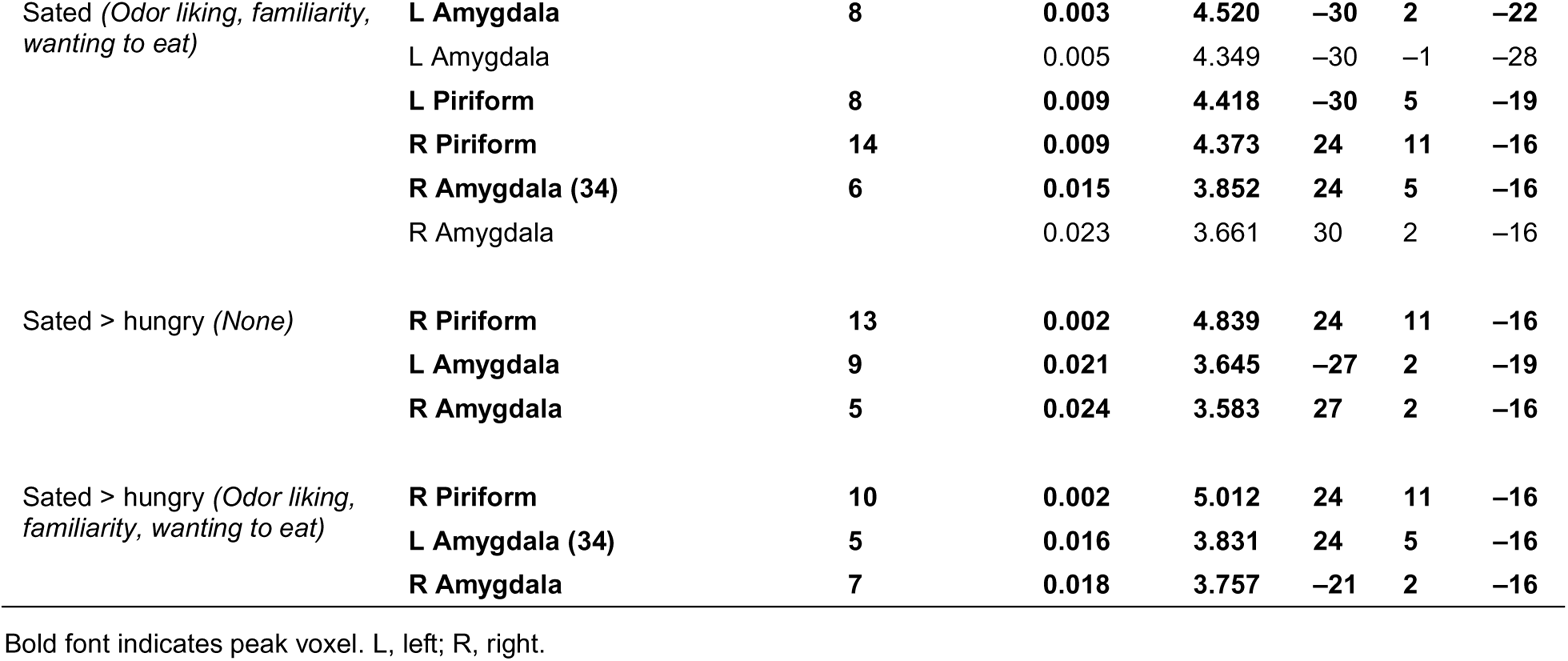
Regions with significant decoding accuracy for food versus nonfood odors.

We next tested the influence of internal state on average odor-evoked brain signals using traditional univariate analyses. In our prior work, we collapsed across odorant category because preliminary analyses showed no differences between food and nonfood odors. This identified significant BOLD responses to all odors > clean air bilaterally in the piriform cortices, the amygdala, and the orbitofrontal cortices, among other regions (Sun et al., 2016). Here we re-ran the analyses separately for the food and nonfood odors, and, as in our previous (but unreported) preliminary analyses, no main effects or interactions with odor category or internal state on fMRI response were found (**Fig. 2c**; all *p*_FWE_ ≥ 0.123). These findings contrast with the decoding results and indicate that the separability of brain patterns responding to the food versus nonfood odors is not attributable to univariate differences.

#### 3.2.2 Post-meal olfactory decoding is associated with energy intake, but not weight change

Our next aim was to determine if olfactory decoding or univariate BOLD responses to food versus nonfood odors correlates with food intake or weight change. Individual differences in the SVM decoding accuracies in the hungry state were unrelated to post-scan *ad libitum* intake adjusted for BMI, sex, age, and subjective satiation (all *p*_FWE_ ≥ 0.340). However, we found that the SVM decoding accuracies in the left amygdala (**Fig. 3a**), parahippocampal gyrus, fusiform gyrus (**Fig. 3b**), cerebellum, middle cingulate gyrus (**Fig. 3c**), and precentral gyrus were positively associated with food intake when sated (**Table 4**). The left piriform cortex and right amygdala also emerged from these analyses after SVC. By contrast, univariate BOLD responses to the food > nonfood odors were unrelated to food intake regardless of internal state (all *p*_FWE_ ≥ 0.100). Decoding accuracies were also unrelated to any odor perceptual rating (all *p*_FWE_ ≥ 0.106), odor dilutions (all *p*_FWE_ ≥ 0.553), composite head motion in the scanner (all *p*_FWE_ ≥ 0.325), sniff vigor (all *p*_FWE_ ≥ 0.488), sniff volume (all *p*_FWE_ ≥ 0.654), or dietary restraint (all *p*_FWE_ ≥ 0.517) in either internal state. Additionally, decoding accuracies were not associated with average total ghrelin when hungry (all *p*_FWE_ ≥ 0.285) or with peak ghrelin excursion when sated (all *p*_FWE_ ≥ 0.547).

**Figure 3.**
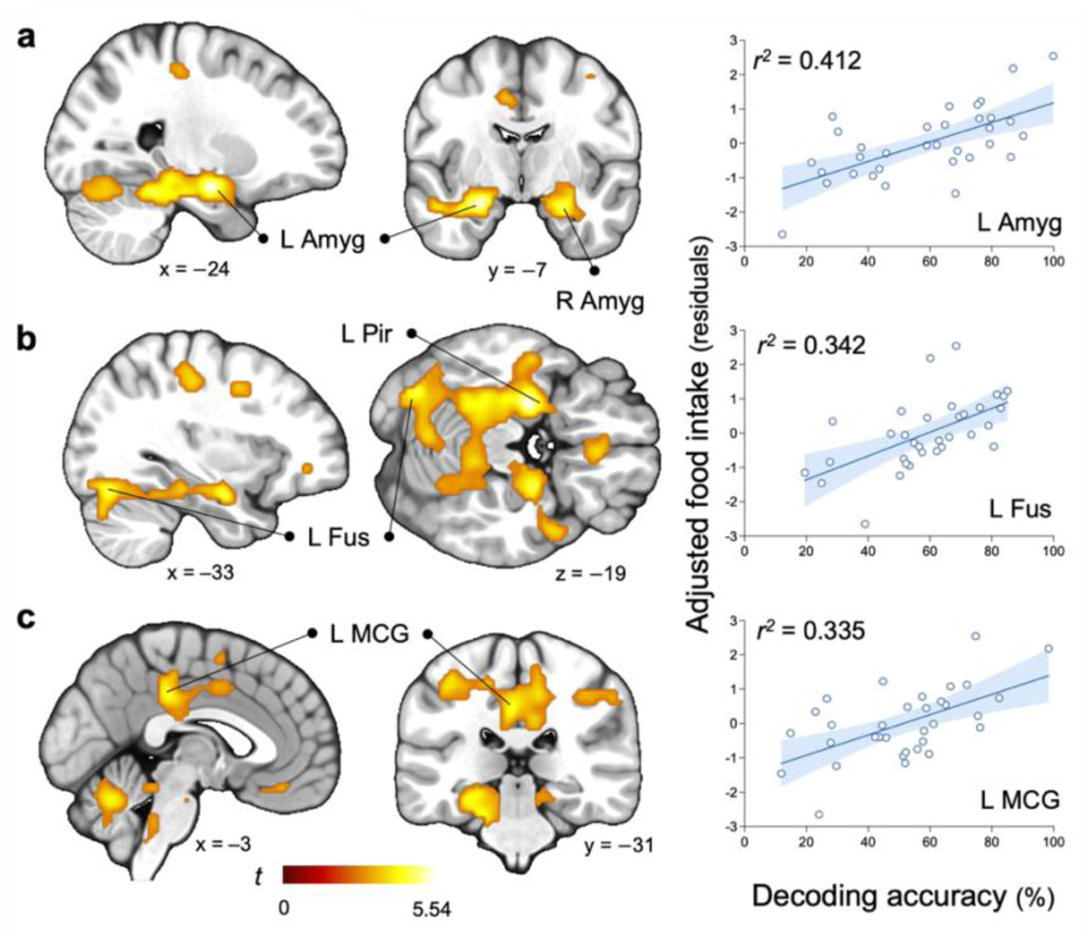
Olfactory decoding is positively associated food intake in the sated state. Fitted scatter plots depict whole-brain-corrected positive associations between post-meal SVM decoding accuracy and adjusted food intake in the left **(a)** amygdala, **(b)** fusiform gyrus, and **(c)** middle cingulate gyrus. Significant effects also emerged in the left piriform cortex and right amygdala from the ROI analyses. Brain sections are shown as in Figure 2. Decoding accuracies were extracted from the peak voxels of the regression with food intake that included covariates for BMI, sex, age, and subjective satiation. The residuals of food intake after adjusting for these factors are plotted with the *r^2^* coefficient values for illustrative purposes. Each data point refers to a single participant and shading indicates the 95% confidence interval. L, left; R, right; Amyg, amygdala; Fus, fusiform gyrus; MCG, middle cingulate gyrus; Pir, piriform.

**Table 4.** Significant associations between decoding accuracy and ad libitum food intake.

| **WHOLE-BRAIN ANALYSES** | | Size (Voxels) | $p_{\text{FWE-cluster}}$ | $t$ value | MNI | | |
| --- | --- | --- | --- | --- | --- | --- | --- |
| Condition (Covariates) | Label (Brodmann Area) |  |  |  | x | y | z |
| Sated (*BMI, sex, age, satiation index*) | **L Amygdala** | **357** | **0.0002** | **5.536** | **-24** | **-7** | **-19** |
|  | L Cerebellum: Anterior lobe culmen |  |  | 4.640 | -24 | -40 | -22 |
|  | L Parahippocampal gyrus |  |  | 4.515 | -30 | -31 | -16 |
|  | **L Fusiform gyrus (19)** | **137** | **0.019** | **4.772** | **-33** | **-73** | **-19** |
|  | L Cerebellum: Posterior lobe declive |  |  | 4.700 | -6 | -64 | -25 |
|  | L Cerebellum: Posterior lobe crus 1 |  |  | 3.720 | -33 | -79 | -31 |
|  | **L Middle cingulate gyrus (31)** | **136** | **0.020** | **4.562** | **-3** | **-31** | **38** |
|  | L Middle cingulate gyrus (24) |  |  | 4.241 | -9 | -13 | 41 |
|  | R Precentral gyrus |  |  | 3.924 | 12 | -31 | 47 |
| **ROI ANALYSES** | | Size (Voxels) | $p_{\text{FWE}}$ | $t$ value | MNI | | |
| Condition (Covariates) | Label (Brodmann Area) |  |  |  | x | y | z |
| Sated (*BMI, sex, age, satiation index*) | **L Amygdala** | **47** | **0.0003** | **5.536** | **-24** | **-7** | **-19** |
|  | **R Amygdala** | **18** | **0.002** | **4.806** | **24** | **-4** | **-19** |
|  | **L Piriform** | **7** | **0.015** | **4.213** | **-27** | **-1** | **-19** |

Bold font indicates peak voxel. L, left; R, right.

There were no significant associations between ΔBMI and olfactory decoding or univariate response when participants were hungry or sated (all *p*_FWE_ ≥ 0.583). An interaction of Taq1A genotype on the relationship between amygdala reactivity to food odors when hungry and weight change has been reported in this dataset (Sun et al., 2015). However, we did not explore the role of the Taq1A polymorphism in the current study due to power limitations with our multivariate techniques.

#### 3.2.3 Amygdala-cerebellum connectivity is associated with olfactory decoding

As a follow-up, we examined whether the extent to which odor category modulates functional connectivity with the amygdala is related to olfactory decoding or food intake. A PPI analysis revealed a significant positive association between food > nonfood connectivity and peak amygdalar decoding accuracy in the right cerebellum (**Fig. 4**; **Table 5**). Connectivity with the left cerebellum also approached statistical significance (*t*_(32)_ = 4.424, *p*_FWE-cluster_ = 0.084, size = 36 voxels, x = –21, y = –82, z = −34). This suggests that individuals with better olfactory decoding in the amygdala show enhanced functional coupling with the cerebellum for food relative to nonfood odors. No significant effects were observed for the relationship between amygdalar connectivity and food intake (all *p*_FWE_ ≥ 0.403).

**Figure 4.**
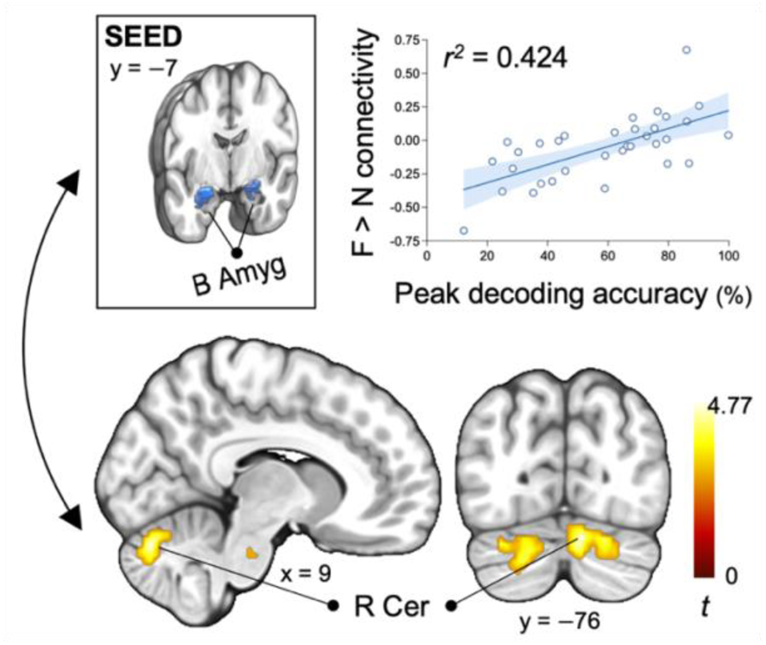
Amygdala-cerebellum connectivity from a PPI analysis is associated with fMRI odor category decoding. Post-meal peak decoding accuracy in the amygdala (x = –24, y = –7, z = –19) positively correlated with functional connectivity differences while sniffing food > nonfood odors between the bilateral amygdala seed and the right cerebellum (crus 1 extending to crus 2 and the declive). Brain sections and scatter plots are shown as in Figures 2 and 3, respectively. Connectivity values were extracted from the peak voxel of the cerebellum (x = 9, y = –76, z = –25) for illustrative purposes. B, bilateral; R, right; Amyg, amygdala; Cer, cerebellum; F, food odors; N, nonfood odors.

**Table 5.**
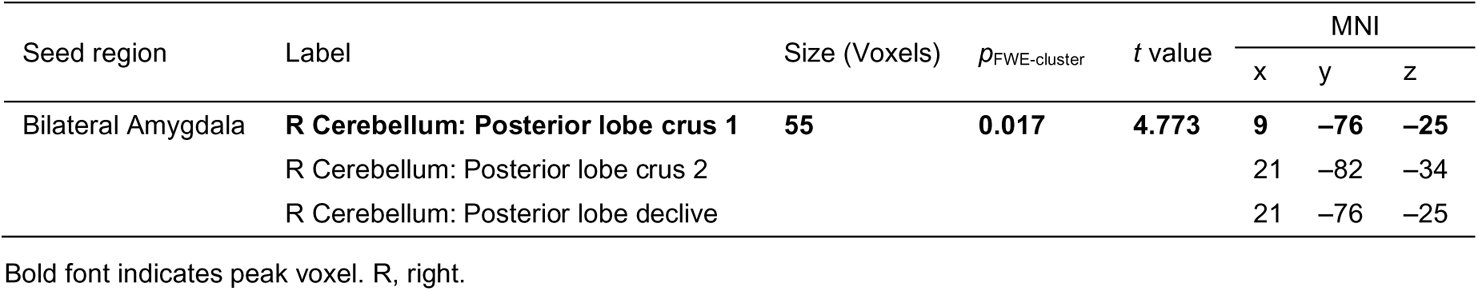
Differential connectivity with the amygdala as a function of decoding accuracy in the sated state.

## 4. Discussion

Controversy exists over the role of olfaction in eating behavior and body weight regulation. Here we used fMRI to test for the existence of relationships between odor category decoding in the brain and (1) internal state, (2) food intake, and (3) weight change over one year. No significant differences for food versus nonfood odors were observed when using traditional univariate comparisons of odor-evoked brain responses. However, decoding accuracies in the piriform primary olfactory cortex and in the amygdala were influenced by internal state and positively associated with *ad libitum* food intake. Odor category decoding accuracies in the fusiform and cingulate cortices were also associated with *ad libitum* intake. By contrast, we found no relationships between differential univariate response or decoding accuracy for food versus nonfood odors and weight change. Taken together, these results (1) demonstrate that internal state influences odor category decoding in the primary olfactory cortex and amygdala, and (2) provide evidence for a positive relationship between decoding accuracy and immediate food intake.

### 4.1 Internal state influences odor category decoding

Prior work has established that odor quality is encoded in patterns of neural responses (Stettler and Axel, 2009) that can be captured in humans using MVPA of odor-evoked fMRI activity (Howard et al., 2009). MVPA can also separate distinct patterns for odorant categories, such as food versus nonfood odors (Bhutani et al., 2019; Howard and Kahnt, 2017; Qu et al., 2016; Shanahan et al., 2021). In the current study, we used MVPA to test the ability of a SVM to decode food versus nonfood odors and demonstrated that its accuracy is significantly greater in the primary olfactory cortex and amygdala in the sated compared to hungry state. This is in line with preclinical findings showing that the neural discrimination of odors in the mouse olfactory bulb decreases with prolonged fasting (Wu et al., 2020). However, it conflicts with a recent fMRI study assessing the effect of energy consumption on food versus nonfood olfactory decoding (Shanahan et al., 2021). Specifically, this study showed that eating a food to satiety reduced the separability of fMRI activity patterns for meal-related food odors versus nonfood odors (e.g., the decoding of pizza versus pine odors after *ad libitum* pizza intake) in olfactory and limbic regions. By contrast, there was no effect of satiation on the discriminability of non-meal-related food odors from nonfood odors (e.g., the decoding of cinnamon bun versus pine odors after *ad libitum* pizza intake). Parallel shifts were also observed in participant ratings of the odor “food-like” qualities, which decreased pre- to post-meal for the meal-matched, but not for the unmatched, food odors. These findings suggest that odor edibility played a role in the impact of satiety on olfactory decoding.

In the current study, the meal was designed to be unrelated to the food odors, which were rated as more liked and desirable to eat than the nonfood odors regardless of internal state. Thus, the improvement in odor category decoding when sated versus hungry is unlikely associated with changes in value or edibility. It also cannot be attributed to differences in intensity, as the odors were all adjusted to be perceived as equally intense. When we further accounted for differences in liking, wanting to eat, and familiarity between the food and nonfood odors in our model, we observed greater decoding accuracy post-meal not only in the piriform and amygdala but also in the midbrain. We speculate that one explanation for the effect of internal state in our study might be that in a state of hunger, a greater number of olfactory percepts are treated as potential food sources within central brain circuits. If so, then the lilac and honeysuckle nonfood odors may also be coded as potential nutrient sources, thereby lowering the accuracy with which the SVM could distinguish these nonfood odors from food odors when hungry. Consequently, in both studies, the key factor determining pattern separation would be whether odors represent a potential new source of energy.

### 4.2 Olfactory decoding is associated with food intake

Accumulating research supports the existence of an association between the olfactory system and the regulation of energy balance, but the nature of the association is unclear. Prior research has reported a positive (Hubert et al., 1980; Schiffman, 1998; Stafford and Whittle, 2015; Riera et al., 2017; Han et al., 2021), negative (Mattes et al., 1990; Mattes and Cowart, 1994; Richardson et al., 2004; Aschenbrenner et al., 2008; Patel et al., 2015; Sun et al., 2015; Han et al., 2020; Temmel et al., 2002), or lack of (Boone et al., 2021; Poessel et al., 2021) relationship between olfaction and obesity risk. Here we demonstrated that the decoding accuracies of food versus nonfood odors in the piriform primary olfactory cortex, fusiform cortex, anterior cingulate cortex, and the amygdala are positively associated with post-scan *ad libitum* energy intake in the sated state. This suggests that better discriminability of these odor category patterns is associated with greater food cue reactivity in the absence of hunger, which is consistent with prior findings demonstrating that food aromas can promote eating (Fedoroff et al. 1997; Fedoroff et al. 2003; Gaillet et al. 2013; Proserpio et al. 2017, 2019), especially in a sated state (Weingarten, 1983; Birch et al., 2003; Goldschmidt et al., 2017).

Positive relationships have also been reported between neural responses to flavors (e.g., milkshakes) and food intake (Burger and Stice, 2013; Nolan-Poupart et al., 2013), but this has not been described for orthonasal odor presentations. This distinction between orthonasal and retronasal olfaction is important because orthonasally sensed odors provide information about the availability of foods and therefore contribute to decisions about food acquisition, whereas retronasal olfaction is the key sensory component of flavor and occurs when the food is already being consumed (Rozin, 1982; Small and Prescott, 2005). Consequently, retronasal olfaction contributes to judgements of acceptability, rather than accessibility. The current finding, therefore, provides a unique link between olfactory decoding and decisions related to food acquisition. In addition, the specificity of our finding to olfactory decoding and not average univariate activation further suggests that orthonasal olfactory processing may guide ingestive behavior through fine alterations in the distributed neural representations of different odor categories rather than large-scale amplification or dampening of the regional brain signal.

### 4.3 Eating and odor decoding beyond olfactory cortex

A growing body of evidence suggests that the fusiform and cingulate cortices convey information relevant to the nutritional properties of foods. For example, the fusiform and middle cingulate gyri are preferentially activated by pictures of foods versus nonfood objects (van der Laan et al., 2011; Murdaugh et al., 2012), and the magnitude of fusiform response to images of fatty foods is negatively associated with participant estimates of their energy densities (DiFeliceantonio et al., 2018). In contrast, a positive relationship is observed for equally liked pictures of foods containing primarily carbohydrate, which participants are worse at estimating, suggesting that it is the accuracy of nutritional information rather than hedonic coding or value that is being represented (DiFeliceantonio et al., 2018). Accordingly, MVPA has revealed that distinct patterns in the fusiform gyri respond to both the actual tastes of gustatory stimuli and the inferred tastes of images depicting sweet, savory, and salty foods (Avery et al., 2021). Responses to food pictures in the fusiform and cingulate cortices are also sensitive to internal state, energy content, body weight status, and sex (LaBar et al., 2001; Siep et al., 2009; Toepel et al., 2009; Frank et al., 2010; Geliebter et al., 2013; Chao et al., 2017; Yeung, 2018; Mengotti et al., 2019). They are further linked with risk for overeating since responses to high-calorie food images in both areas negatively predict weight loss and weight loss maintenance (Murdaugh et al., 2012). The results of the current study add to this literature by showing that greater decoding accuracies of food versus nonfood odor categories in the fusiform and cingulate cortices are associated with greater *ad libitum* intake specifically in the sated state. Collectively, these findings suggest that these regions encode nutritional information from environmental and internal sources and that variation in the fidelity of this coding is associated with eating behavior.

There is also emerging evidence for cerebellar involvement in food intake. Human neuroimaging studies have reported cerebellar responses to food-related stimuli (Huerta et al., 2014) that are blunted by satiation (Tataranni et al., 1999; Small et al., 2001) and in hyperphagic individuals with Prader-Willi syndrome (Blanco-Hinojo et al., 2019; Low et al., 2021). There is also a population of neurons in the deep cerebellar nuclei that are proposed to act as a satiation network by modulating the dopaminergic system toward reducing food intake in mice (Low et al., 2021). In the current study, we found that better olfactory decoding in the amygdala when sated is positively associated with stronger functional coupling between the amygdala and the right cerebellum crus I/II and declive in response to food relative to nonfood odors. While these regions do not correspond to the deep cerebellar nuclei, they do overlap with prior neuroimaging reports of cerebellar reactivity while viewing pictures of high versus low-calorie foods (Killgore et al., 2003) and smelling (Sobel et al., 1998) and discriminating (Savic et al., 2000) odor qualities.

### 4.4 Caveats and limitations

Participants rated the food odors as more liked, wanted, and familiar than the nonfood odors in the hungry and sated states (Table 1). Although these perceptual ratings were consistently used as covariates in our models, we cannot definitively rule out the possibility that decoding accuracy was influenced not only by category, but also by these variables. However, we believe this is unlikely since the decoding accuracies, but not perceptual ratings, were influenced by internal state. Another limitation is the small sample size. Although multivariate methods tend to produce more reproducible brain-wide association effects than univariate techniques (Marek et al., 2022), it is possible that the lack of association between decoding accuracy and weight change reflects insufficient sensitivity rather than a true null effect. Finally, there were insufficient presentations of each odor quality during fMRI scanning to perform MVPA of all odor qualities here (e.g., lilac versus honeysuckle). Future work should test if the relationship between olfactory decoding and food intake is specific to food versus nonfood odors or extends to other odor categories and qualities.

### 4.5 Conclusions

We demonstrate that the decoding accuracies of fMRI activity patterns for food versus nonfood odor categories in the primary olfactory cortex and the amygdala are superior in the sated state. Decoding accuracies in these regions and in the fusiform and cingulate cortices are also positively associated with *ad libitum* energy consumption. These results are consistent with prior studies showing that internal state influences olfactory decoding and with work suggesting that enhanced olfactory function may be associated with increased food intake.

## Data availability

The data are available upon request from the corresponding authors.

## Acknowledgements

We would like to thank the additional individuals that contributed to the original studies that this retrospective analysis is based upon, including Nils Kroemer, Maria G. Veldhuizen, Amanda E. Babbs, Ivan E. de Araujo, Darren R. Gitelman, Robert S. Sherwin, and Rajita Sinha.

## Authors’ contributions

Emily E. Perszyk, Conceptualization, Methodology, Formal analysis, Data curation, Writing – original draft preparation, Visualization, Funding acquisition; Xue S. Davis, Conceptualization, Methodology, Formal analysis, Investigation, Data curation, Writing – review and editing; Dana M. Small, Conceptualization, Resources, Writing – original draft preparation, Supervision, Funding acquisition. All authors have read and approved the final article.

## References

Aimé, P., Duchamp-Viret, P., Chaput, M.A., Savigner, A., Mahfouz, M., Julliard, A.K., 2007. Fasting increases and satiation decreases olfactory detection for a neutral odor in rats. Behavioural Brain Research 179, 258–264. https://doi.org/10.1016/j.bbr.2007.02.012

Aschenbrenner, K., Hummel, C., Teszmer, K., Krone, F., Ishimaru, T., Seo, H.-S., Hummel, T., 2008. The Influence of Olfactory Loss on Dietary Behaviors. The Laryngoscope 118, 135–144. https://doi.org/10.1097/MLG.0b013e318155a4b9

Avery, J.A., Liu, A.G., Ingeholm, J.E., Gotts, S.J., Martin, A., 2021. Viewing images of foods evokes taste quality-specific activity in gustatory insular cortex. PNAS 118. https://doi.org/10.1073/pnas.2010932118

Bao, X., Raguet, L.L., Cole, S.M., Howard, J.D., Gottfried, J.A., 2016. The role of piriform associative connections in odor categorization. eLife 5, e13732. https://doi.org/10.7554/eLife.13732

Bartoshuk, L.M., Duffy, V.B., Green, B.G., Hoffman, H.J., Ko, C.-W., Lucchina, L.A., Marks, L.E., Snyder, D.J., Weiffenbach, J.M., 2004. Valid across-group comparisons with labeled scales: the gLMS versus magnitude matching. Physiology & Behavior, Festschrift in Honor of Gerard P. Smith 82, 109–114. https://doi.org/10.1016/j.physbeh.2004.02.033

Betley, J.N., Xu, S., Cao, Z.F.H., Gong, R., Magnus, C.J., Yu, Y., Sternson, S.M., 2015. Neurons for hunger and thirst transmit a negative-valence teaching signal. Nature 521, 180–185. https://doi.org/10.1038/nature14416

Bhutani, S., Howard, J.D., Reynolds, R., Zee, P.C., Gottfried, J., Kahnt, T., 2019. Olfactory connectivity mediates sleep-dependent food choices in humans. eLife 8. https://doi.org/10.7554/eLife.49053

Birch, L.L., Fisher, J.O., Davison, K.K., 2003. Learning to overeat: maternal use of restrictive feeding practices promotes girls’ eating in the absence of hunger. Am J Clin Nutr 78, 215–220. https://doi.org/10.1093/ajcn/78.2.215

Blanco-Hinojo, L., Pujol, J., Esteba-Castillo, S., Martínez-Vilavella, G., Giménez-Palop, O., Gabau, E., Casamitjana, L., Deus, J., Novell, R., Caixàs, A., 2019. Lack of response to disgusting food in the hypothalamus and related structures in Prader Willi syndrome. NeuroImage: Clinical 21, 101662. https://doi.org/10.1016/j.nicl.2019.101662

Boone, M.H., Liang-Guallpa, J., Krashes, M.J., 2021. Examining the role of olfaction in dietary choice. Cell Reports 34, 108755. https://doi.org/10.1016/j.celrep.2021.108755

Boswell, R.G., Kober, H., 2016. Food cue reactivity and craving predict eating and weight gain: a meta-analytic review. Obes Rev 17, 159–177. https://doi.org/10.1111/obr.12354

Burger, K.S., Stice, E., 2013. Elevated energy intake is correlated with hyperresponsivity in attentional, gustatory, and reward brain regions while anticipating palatable food receipt. The American Journal of Clinical Nutrition 97, 1188–1194. https://doi.org/10.3945/ajcn.112.055285

Cameron, J.D., Goldfield, G.S., Doucet, É., 2012. Fasting for 24h improves nasal chemosensory performance and food palatability in a related manner. Appetite 58, 978–981. https://doi.org/10.1016/j.appet.2012.02.050

Chang, C.-C., Lin, C.-J., 2011. LIBSVM: A library for support vector machines. ACM Trans. Intell. Syst. Technol. 2, 27:1–27:27. https://doi.org/10.1145/1961189.1961199

Chao, A.M., Loughead, J., Bakizada, Z.M., Hopkins, C.M., Geliebter, A., Gur, R.C., Wadden, T.A., 2017. Sex/gender differences in neural correlates of food stimuli: a systematic review of functional neuroimaging studies. Obesity Reviews 18, 687–699. https://doi.org/10.1111/obr.12527

Chen, Y., Lin, Y.-C., Kuo, T.-W., Knight, Z.A., 2015. Sensory Detection of Food Rapidly Modulates Arcuate Feeding Circuits. Cell 160, 829–841. https://doi.org/10.1016/j.cell.2015.01.033

Colbert, H.A., Bargmann, C.I., 1997. Environmental signals modulate olfactory acuity, discrimination, and memory in Caenorhabditis elegans. Learn. Mem. 4, 179–191. https://doi.org/10.1101/lm.4.2.179

DiFeliceantonio, A.G., Coppin, G., Rigoux, L., Edwin Thanarajah, S., Dagher, A., Tittgemeyer, M., Small, D.M., 2018. Supra-Additive Effects of Combining Fat and Carbohydrate on Food Reward. Cell Metabolism 28, 33–44.e3. https://doi.org/10.1016/j.cmet.2018.05.018

Elmquist, J.K., Bjørbæk, C., Ahima, R.S., Flier, J.S., Saper, C.B., 1998. Distributions of leptin receptor mRNA isoforms in the rat brain. Journal of Comparative Neurology 395, 535–547. https://doi.org/10.1002/(SICI)1096-9861(19980615)395:4<535::AID-CNE9>3.0.CO;2-2

Fadool, D.A., Tucker, K., Perkins, R., Fasciani, G., Thompson, R.N., Parsons, A.D., Overton, J.M., Koni, P.A., Flavell, R.A., Kaczmarek, L.K., 2004. Kv1.3 Channel Gene-Targeted Deletion Produces “Super-Smeller Mice” with Altered Glomeruli, Interacting Scaffolding Proteins, and Biophysics. Neuron 41, 389–404. https://doi.org/10.1016/S0896-6273(03)00844-4

Fedoroff, I., Polivy, J., Peter Herman, C., 2003. The specificity of restrained versus unrestrained eaters’ responses to food cues: general desire to eat, or craving for the cued food? Appetite 41, 7–13. https://doi.org/10.1016/S0195-6663(03)00026-6

Fedoroff, I.D.C., Polivy, J., Herman, C.P., 1997. The Effect of Pre-exposure to Food Cues on the Eating Behavior of Restrained and Unrestrained Eaters. Appetite 28, 33–47. https://doi.org/10.1006/appe.1996.0057

Feldman, M., Richardson, C.T., 1986. Role of thought, sight, smell, and taste of food in the cephalic phase of gastric acid secretion in humans. Gastroenterology 90, 428–433. https://doi.org/10.1016/0016-5085(86)90943-1

Frank, S., Laharnar, N., Kullmann, S., Veit, R., Canova, C., Hegner, Y.L., Fritsche, A., Preissl, H., 2010. Processing of food pictures: Influence of hunger, gender and calorie content. Brain Research, Neural Mechanisms of Ingestive Behaviour and Obesity 1350, 159–166. https://doi.org/10.1016/j.brainres.2010.04.030

Friston, K.J., Buechel, C., Fink, G.R., Morris, J., Rolls, E., Dolan, R.J., 1997. Psychophysiological and Modulatory Interactions in Neuroimaging. NeuroImage 6, 218–229. https://doi.org/10.1006/nimg.1997.0291

Gaillet, M., Sulmont-Rossé, C., Issanchou, S., Chabanet, C., Chambaron, S., 2013. Priming effects of an olfactory food cue on subsequent food-related behaviour. Food Quality and Preference 30, 274–281. https://doi.org/10.1016/j.foodqual.2013.06.008

Gall, C., Seroogy, K.B., Brecha, N., 1986. Distribution of VIP- and NPY-like immunoreactivities in rat main olfactory bulb. Brain Research 374, 389–394. https://doi.org/10.1016/0006-8993(86)90436-1

Geliebter, A., Pantazatos, S.P., McOuatt, H., Puma, L., Gibson, C.D., Atalayer, D., 2013. Sex-based fMRI differences in obese humans in response to high vs. low energy food cues. Behavioural Brain Research 243, 91–96. https://doi.org/10.1016/j.bbr.2012.12.023

Gitelman, D.R., Penny, W.D., Ashburner, J., Friston, K.J., 2003. Modeling regional and psychophysiologic interactions in fMRI: the importance of hemodynamic deconvolution. NeuroImage 19, 200–207. https://doi.org/10.1016/S1053-8119(03)00058-2

Goldschmidt, A.B., Crosby, R.D., Cao, L., Pearson, C.M., Utzinger, L.M., Pacanowski, C.R., Mason, T.B., Berner, L.A., Engel, S.G., Wonderlich, S.A., Peterson, C.B., 2017. Contextual factors associated with eating in the absence of hunger among adults with obesity. Eating Behaviors 26, 33–39. https://doi.org/10.1016/j.eatbeh.2017.01.005

Green, B.G., Dalton, P., Cowart, B., Shaffer, G., Rankin, K., Higgins, J., 1996. Evaluating the ‘Labeled Magnitude Scale’ for Measuring Sensations of Taste and Smell. Chemical Senses 21, 323–334. https://doi.org/10.1093/chemse/21.3.323

Green, B.G., Shaffer, G.S., Gilmore, M.M., 1993. Derivation and evaluation of a semantic scale of oral sensation magnitude with apparent ratio properties. Chemical Senses 18, 683–702. https://doi.org/10.1093/chemse/18.6.683

Han, P., Chen, H., Hummel, T., 2020. Brain Responses to Food Odors Associated With BMI Change at 2-Year Follow-Up. Frontiers in Human Neuroscience 14, 402. https://doi.org/10.3389/fnhum.2020.574148

Han, P., Roitzsch, C., Horstmann, A., Pössel, M., Hummel, T., 2021. Increased Brain Reward Responsivity to Food-Related Odors in Obesity. Obesity 29, 1138–1145. https://doi.org/10.1002/oby.23170

Hebart, M.N., Görgen, K., Haynes, J.-D., 2015. The Decoding Toolbox (TDT): a versatile software package for multivariate analyses of functional imaging data. Front. Neuroinform. 8. https://doi.org/10.3389/fninf.2014.00088

Hill, J.M., Lesniak, M.A., Pert, C.B., Roth, J., 1986. Autoradiographic localization of insulin receptors in rat brain: Prominence in olfactory and limbic areas. Neuroscience 17, 1127–1138. https://doi.org/10.1016/0306-4522(86)90082-5

Howard, J.D., Kahnt, T., 2017. Identity-Specific Reward Representations in Orbitofrontal Cortex Are Modulated by Selective Devaluation. J. Neurosci. 37, 2627–2638. https://doi.org/10.1523/JNEUROSCI.3473-16.2017

Howard, J.D., Plailly, J., Grueschow, M., Haynes, J.-D., Gottfried, J.A., 2009. Odor quality coding and categorization in human posterior piriform cortex. Nature Neuroscience 12, 932–938. https://doi.org/10.1038/nn.2324

Hubert, H.B., Fabsitz, R.R., Feinleib, M., Brown, K.S., 1980. Olfactory sensitivity in humans: genetic versus environmental control. Science 208, 607–609. https://doi.org/10.1126/science.7189296

Huerta, C.I., Sarkar, P.R., Duong, T.Q., Laird, A.R., Fox, P.T., 2014. Neural bases of food perception: Coordinate-based meta-analyses of neuroimaging studies in multiple modalities. Obesity 22, 1439–1446. https://doi.org/10.1002/oby.20659

Johnson, B.N., Sobel, N., 2007. Methods for building an olfactometer with known concentration outcomes. Journal of Neuroscience Methods 160, 231–245. https://doi.org/10.1016/j.jneumeth.2006.09.008

Julliard, A.K., Chaput, M.A., Apelbaum, A., Aimé, P., Mahfouz, M., Duchamp-Viret, P., 2007. Changes in rat olfactory detection performance induced by orexin and leptin mimicking fasting and satiation. Behavioural Brain Research 183, 123–129. https://doi.org/10.1016/j.bbr.2007.05.033

Killgore, W.D.S., Young, A.D., Femia, L.A., Bogorodzki, P., Rogowska, J., Yurgelun-Todd, D.A., 2003. Cortical and limbic activation during viewing of high- versus low-calorie foods. NeuroImage 19, 1381–1394. https://doi.org/10.1016/S1053-8119(03)00191-5

LaBar, K.S., Gitelman, D.R., Parrish, T.B., Kim, Y.-H., Nobre, A.C., Mesulam, M.-M., 2001. Hunger selectively modulates corticolimbic activation to food stimuli in humans. Behavioral Neuroscience 115, 493–500. https://doi.org/10.1037/0735-7044.115.2.493

Lee, V.M., Linden, R.W.A., 1992. The effect of odours on stimulated parotid salivary flow in humans. Physiology & Behavior 52, 1121–1125. https://doi.org/10.1016/0031-9384(92)90470-M

Lim, J., Wood, A., Green, B.G., 2009. Derivation and Evaluation of a Labeled Hedonic Scale. Chem Senses 34, 739–751. https://doi.org/10.1093/chemse/bjp054

Low, A.Y.T., Goldstein, N., Gaunt, J.R., Huang, K.-P., Zainolabidin, N., Yip, A.K.K., Carty, J.R.E., Choi, J.Y., Miller, A.M., Ho, H.S.T., Lenherr, C., Baltar, N., Azim, E., Sessions, O.M., Ch’ng, T.H., Bruce, A.S., Martin, L.E., Halko, M.A., Brady, R.O., Holsen, L.M., Alhadeff, A.L., Chen, A.I., Betley, J.N., 2021. Reverse-translational identification of a cerebellar satiation network. Nature 1–5. https://doi.org/10.1038/s41586-021-04143-5

Macey, P.M., Macey, K.E., Kumar, R., Harper, R.M., 2004. A method for removal of global effects from fMRI time series. NeuroImage 22, 360–366. https://doi.org/10.1016/j.neuroimage.2003.12.042

Marek, S., Tervo-Clemmens, B., Calabro, F.J., Montez, D.F., Kay, B.P., Hatoum, A.S., Donohue, M.R., Foran, W., Miller, R.L., Hendrickson, T.J., Malone, S.M., Kandala, S., Feczko, E., Miranda-Dominguez, O., Graham, A.M., Earl, E.A., Perrone, A.J., Cordova, M., Doyle, O., Moore, L.A., Conan, G.M., Uriarte, J., Snider, K., Lynch, B.J., Wilgenbusch, J.C., Pengo, T., Tam, A., Chen, J., Newbold, D.J., Zheng, A., Seider, N.A., Van, A.N., Metoki, A., Chauvin, R.J., Laumann, T.O., Greene, D.J., Petersen, S.E., Garavan, H., Thompson, W.K., Nichols, T.E., Yeo, B.T.T., Barch, D.M., Luna, B., Fair, D.A., Dosenbach, N.U.F., 2022. Reproducible brain-wide association studies require thousands of individuals. Nature 603, 654–660. https://doi.org/10.1038/s41586-022-04492-9

Martí-Henneberg, C., Capdevila, F., Arija, V., Pérez, S., Cucó, G., Vizmanos, B., Fernández-Ballart, J., 1999. Energy density of the diet, food volume and energy intake by age and sex in a healthy population. Eur J Clin Nutr 53, 421–428. https://doi.org/10.1038/sj.ejcn.1600770

Mattes, R.D., Cowart, B.J., 1994. Dietary assessment of patients with chemosensotyr disorders. Journal of the American Dietetic Association 94, 50–56. https://doi.org/10.1016/0002-8223(94)92041-9

Mattes, R.D., Cowart, B.J., Schiavo, M.A., Arnold, C., Garrison, B., Kare, M.R., Lowry, L.D., 1990. Dietary evaluation of patients with smell and/or taste disorders. The American Journal of Clinical Nutrition 51, 233–240. https://doi.org/10.1093/ajcn/51.2.233

McIntyre, J.C., Thiebaud, N., McGann, J.P., Komiyama, T., Rothermel, M., 2017. Neuromodulation in Chemosensory Pathways. Chemical Senses 42, 375–379. https://doi.org/10.1093/chemse/bjx014

McLaren, D.G., Ries, M.L., Xu, G., Johnson, S.C., 2012. A generalized form of context-dependent psychophysiological interactions (gPPI): A comparison to standard approaches. NeuroImage 61, 1277–1286. https://doi.org/10.1016/j.neuroimage.2012.03.068

Mengotti, P., Foroni, F., Rumiati, R.I., 2019. Neural correlates of the energetic value of food during visual processing and response inhibition. NeuroImage 184, 130–139. https://doi.org/10.1016/j.neuroimage.2018.09.017

Murdaugh, D.L., Cox, J.E., Cook, E.W., Weller, R.E., 2012. fMRI reactivity to high-calorie food pictures predicts short- and long-term outcome in a weight-loss program. NeuroImage 59, 2709–2721. https://doi.org/10.1016/j.neuroimage.2011.10.071

Nolan-Poupart, S., Veldhuizen, M.G., Geha, P., Small, D.M., 2013. Midbrain response to milkshake correlates with ad libitum milkshake intake in the absence of hunger. Appetite 60, 168–174. https://doi.org/10.1016/j.appet.2012.09.032

Pager, J., Giachetti, I., Holley, A., Le Magnen, J., 1972. A selective control of olfactory bulb electrical activity in relation to food deprivation and satiety in rats. Physiology & Behavior 9, 573–579. https://doi.org/10.1016/0031-9384(72)90014-5

Palouzier-Paulignan, B., Lacroix, M.-C., Aimé, P., Baly, C., Caillol, M., Congar, P., Julliard, A.K., Tucker, K., Fadool, D.A., 2012. Olfaction Under Metabolic Influences. Chemical Senses 37, 769–797. https://doi.org/10.1093/chemse/bjs059

Patel, Z.M., DelGaudio, J.M., Wise, S.K., 2015. Higher Body Mass Index Is Associated with Subjective Olfactory Dysfunction. Behavioural Neurology 2015, e675635. https://doi.org/10.1155/2015/675635

Poessel, M., Morys, F., Breuer, N., Villringer, A., Hummel, T., Horstmann, A., 2021. Brain response to food odors is not associated with body mass index and obesity-related metabolic health measures. Appetite 105774. https://doi.org/10.1016/j.appet.2021.105774

Proserpio, C., de Graaf, C., Laureati, M., Pagliarini, E., Boesveldt, S., 2017. Impact of ambient odors on food intake, saliva production and appetite ratings. Physiology & Behavior 174, 35–41. https://doi.org/10.1016/j.physbeh.2017.02.042

Proserpio, C., Invitti, C., Boesveldt, S., Pasqualinotto, L., Laureati, M., Cattaneo, C., Pagliarini, E., 2019. Ambient Odor Exposure Affects Food Intake and Sensory Specific Appetite in Obese Women. Frontiers in Psychology 10, 7. https://doi.org/10.3389/fpsyg.2019.00007

Prud’homme, M.J., Lacroix, M.C., Badonnel, K., Gougis, S., Baly, C., Salesse, R., Caillol, M., 2009. Nutritional status modulates behavioural and olfactory bulb Fos responses to isoamyl acetate or food odour in rats: roles of orexins and leptin. Neuroscience 162, 1287–1298. https://doi.org/10.1016/j.neuroscience.2009.05.043

Qu, L.P., Kahnt, T., Cole, S.M., Gottfried, J.A., 2016. De Novo Emergence of Odor Category Representations in the Human Brain. J. Neurosci. 36, 468–478. https://doi.org/10.1523/JNEUROSCI.3248-15.2016

Richardson, B.E., Vander Woude, E.A., Sudan, R., Thompson, J.S., Leopold, D.A., 2004. Altered Olfactory Acuity in the Morbidly Obese. OBES SURG 14, 967–969. https://doi.org/10.1381/0960892041719617

Riera, C.E., Tsaousidou, E., Halloran, J., Follett, P., Hahn, O., Pereira, M.M.A., Ruud, L.E., Alber, J., Tharp, K., Anderson, C.M., Brönneke, H., Hampel, B., Filho, C.D. de M., Stahl, A., Brüning, J.C., Dillin, A., 2017. The Sense of Smell Impacts Metabolic Health and Obesity. Cell Metabolism 26, 198–211.e5. https://doi.org/10.1016/j.cmet.2017.06.015

Rolls, E.T., Huang, C.-C., Lin, C.-P., Feng, J., Joliot, M., 2020. Automated anatomical labelling atlas 3. NeuroImage 206, 116189. https://doi.org/10.1016/j.neuroimage.2019.116189

Root, C.M., Ko, K.I., Jafari, A., Wang, J.W., 2011. Presynaptic Facilitation by Neuropeptide Signaling Mediates Odor-Driven Food Search. Cell 145, 133–144. https://doi.org/10.1016/j.cell.2011.02.008

Rozin, P., 1982. “Taste–smell confusions” and the duality of the olfactory sense. Perception & Psychophysics 31, 397–401. https://doi.org/10.3758/BF03202667

Savic, I., Gulyas, B., Larsson, M., Roland, P., 2000. Olfactory Functions Are Mediated by Parallel and Hierarchical Processing. Neuron 26, 735–745. https://doi.org/10.1016/S0896-6273(00)81209-X

Schiffman, S.S., 1998. Sensory enhancement of foods for the elderly with monosodium glutamate and flavors. Food Reviews International 14, 321–333. https://doi.org/10.1080/87559129809541164

Shanahan, L.K., Bhutani, S., Kahnt, T., 2021. Olfactory perceptual decision-making is biased by motivational state. PLOS Biology 19, e3001374. https://doi.org/10.1371/journal.pbio.3001374

Siep, N., Roefs, A., Roebroeck, A., Havermans, R., Bonte, M.L., Jansen, A., 2009. Hunger is the best spice: An fMRI study of the effects of attention, hunger and calorie content on food reward processing in the amygdala and orbitofrontal cortex. Behavioural Brain Research 198, 149–158. https://doi.org/10.1016/j.bbr.2008.10.035

Small, D.M., Prescott, J., 2005. Odor/taste integration and the perception of flavor. Exp Brain Res 166, 345–357. https://doi.org/10.1007/s00221-005-2376-9

Small, D.M., Veldhuizen, M.G., Felsted, J., Mak, Y.E., McGlone, F., 2008. Separable Substrates for Anticipatory and Consummatory Food Chemosensation. Neuron 57, 786–797. https://doi.org/10.1016/j.neuron.2008.01.021

Small, D.M., Zatorre, R.J., Dagher, A., Evans, A.C., Jones-Gotman, M., 2001. Changes in brain activity related to eating chocolate: From pleasure to aversion. Brain 124, 1720–1733. https://doi.org/10.1093/brain/124.9.1720

Sobel, N., Prabhakaran, V., Hartley, C.A., Desmond, J.E., Zhao, Z., Glover, G.H., Gabrieli, J.D.E., Sullivan, E.V., 1998. Odorant-Induced and Sniff-Induced Activation in the Cerebellum of the Human. J. Neurosci. 18, 8990–9001. https://doi.org/10.1523/JNEUROSCI.18-21-08990.1998

Stafford, L.D., Whittle, A., 2015. Obese Individuals Have Higher Preference and Sensitivity to Odor of Chocolate. Chem Senses 40, 279–284. https://doi.org/10.1093/chemse/bjv007

Stettler, D.D., Axel, R., 2009. Representations of Odor in the Piriform Cortex. Neuron 63, 854–864. https://doi.org/10.1016/j.neuron.2009.09.005

Stevenson, R.J., 2010. An Initial Evaluation of the Functions of Human Olfaction. Chemical Senses 35, 3–20. https://doi.org/10.1093/chemse/bjp083

Strien, T. van, Frijters, J.E.R., Bergers, G.P.A., Defares, P.B., 1986. The Dutch Eating Behavior Questionnaire (DEBQ) for assessment of restrained, emotional, and external eating behavior. International Journal of Eating Disorders 5, 295–315. https://doi.org/10.1002/1098-108X(198602)5:2<295::AID-EAT2260050209>3.0.CO;2-T

Sun, X., Kroemer, N.B., Veldhuizen, M.G., Babbs, A.E., Araujo, I.E. de, Gitelman, D.R., Sherwin, R.S., Sinha, R., Small, D.M., 2015. Basolateral Amygdala Response to Food Cues in the Absence of Hunger Is Associated with Weight Gain Susceptibility. J. Neurosci. 35, 7964–7976. https://doi.org/10.1523/JNEUROSCI.3884-14.2015

Sun, X., Veldhuizen, M.G., Babbs, A.E., Sinha, R., Small, D.M., 2016. Perceptual and Brain Response to Odors Is Associated with Body Mass Index and Postprandial Total Ghrelin Reactivity to a Meal. Chem Senses 41, 233–248. https://doi.org/10.1093/chemse/bjv081

Sun, X., Veldhuizen, M.G., Wray, A.E., de Araujo, I.E., Sherwin, R.S., Sinha, R., Small, D.M., 2014. The neural signature of satiation is associated with ghrelin response and triglyceride metabolism. Physiology & Behavior, SSIB 2014 136, 63–73. https://doi.org/10.1016/j.physbeh.2014.04.017

Tataranni, P.A., Gautier, J.-F., Chen, K., Uecker, A., Bandy, D., Salbe, A.D., Pratley, R.E., Lawson, M., Reiman, E.M., Ravussin, E., 1999. Neuroanatomical correlates of hunger and satiation in humans using positron emission tomography. PNAS 96, 4569–4574. https://doi.org/10.1073/pnas.96.8.4569

Temmel, A.F.P., Quint, C., Schickinger-Fischer, B., Klimek, L., Stoller, E., Hummel, T., 2002. Characteristics of Olfactory Disorders in Relation to Major Causes of Olfactory Loss. Archives of Otolaryngology–Head & Neck Surgery 128, 635–641. https://doi.org/10.1001/archotol.128.6.635

Thiebaud, N., Llewellyn-Smith, I.J., Gribble, F., Reimann, F., Trapp, S., Fadool, D.A., 2016. The incretin hormone glucagon-like peptide 1 increases mitral cell excitability by decreasing conductance of a voltage-dependent potassium channel. The Journal of Physiology 594, 2607–2628. https://doi.org/10.1113/JP272322

Toepel, U., Knebel, J.-F., Hudry, J., le Coutre, J., Murray, M.M., 2009. The brain tracks the energetic value in food images. NeuroImage 44, 967–974. https://doi.org/10.1016/j.neuroimage.2008.10.005

Tong, J., Mannea, E., Aimé, P., Pfluger, P.T., Yi, C.-X., Castaneda, T.R., Davis, H.W., Ren, X., Pixley, S., Benoit, S., Julliard, K., Woods, S.C., Horvath, T.L., Sleeman, M.M., D’Alessio, D., Obici, S., Frank, R., Tschöp, M.H., 2011. Ghrelin Enhances Olfactory Sensitivity and Exploratory Sniffing in Rodents and Humans. J. Neurosci. 31, 5841–5846. https://doi.org/10.1523/JNEUROSCI.5680-10.2011

van der Laan, L.N., de Ridder, D.T.D., Viergever, M.A., Smeets, P.A.M., 2011. The first taste is always with the eyes: A meta-analysis on the neural correlates of processing visual food cues. NeuroImage 55, 296–303. https://doi.org/10.1016/j.neuroimage.2010.11.055

Weingarten, H.P., 1983. Conditioned cues elicit feeding in sated rats: a role for learning in meal initiation. Science 220, 431–433. https://doi.org/10.1126/science.6836286

Wu, J., Liu, P., Chen, F., Ge, L., Lu, Y., Li, A., 2020. Excitability of Neural Activity is Enhanced, but Neural Discrimination of Odors is Slightly Decreased, in the Olfactory Bulb of Fasted Mice. Genes 11, 433. https://doi.org/10.3390/genes11040433

Xu, J., Koni, P.A., Wang, P., Li, G., Kaczmarek, L., Wu, Y., Li, Y., Flavell, R.A., Desir, G.V., 2003. The voltage-gated potassium channel Kv1.3 regulates energy homeostasis and body weight. Human Molecular Genetics 12, 551–559. https://doi.org/10.1093/hmg/ddg049

Yeomans, M.R., 2006. Olfactory influences on appetite and satiety in humans. Physiology & Behavior 87, 800–804. https://doi.org/10.1016/j.physbeh.2006.01.029

Yeung, A.W.K., 2018. Sex differences in brain responses to food stimuli: a meta-analysis on neuroimaging studies. Obesity Reviews 19, 1110–1115. https://doi.org/10.1111/obr.12697

Zhou, G., Lane, G., Cooper, S.L., Kahnt, T., Zelano, C., 2019. Characterizing functional pathways of the human olfactory system. eLife 8, e47177. https://doi.org/10.7554/eLife.47177

